# SpecDB: A Relational Database for Archiving Biomolecular NMR Spectra Data

**DOI:** 10.1101/2022.02.12.480212

**Authors:** Keith J. Fraga, Yuanpeng J. Huang, Theresa A. Ramelot, G.V.T. Swapna, Arwin Lashawn Anak Kendary, Ethan Li, Ian Korf, Gaetano T. Montelione

## Abstract

NMR is a valuable experimental tool in the structural biologist’s toolkit to elucidate the structures, functions, and motions of biomolecules. The progress of machine learning, particularly in structural biology, reveals the critical importance of large, diverse, and reliable datasets in developing new methods and understanding in structural biology and science more broadly. Protein NMR research groups produce large amounts of data, and there is renewed interest in organizing this data to train new, sophisticated machine learning architectures to improve biomolecular NMR analysis pipelines. The foundational data type in NMR is the free-induction decay (FID). There are opportunities to build sophisticated machine learning methods to tackle long-standing problems in NMR data processing, resonance assignment, dynamics analysis, and structure determination using NMR FIDs. Our goal in this study is to provide a lightweight, broadly available tool for archiving FID data as it is generated at the spectrometer, and grow a new resource of FID data and associated metadata. This study presents a relational schema for storing and organizing the metadata items that describe an NMR sample and FID data, which we call **<u>Spec</u>**tra **<u>D</u>**ata**<u>b</u>**ase (SpecDB). SpecDB is implemented in SQLite and includes a Python software library providing a command-line application to create, organize, query, backup, share, and maintain the database. This set of software tools and database schema allow users to store, organize, share, and learn from NMR time domain data. SpecDB is freely available under an open source license at https://github.rpi.edu/RPIBioinformatics/SpecDB.

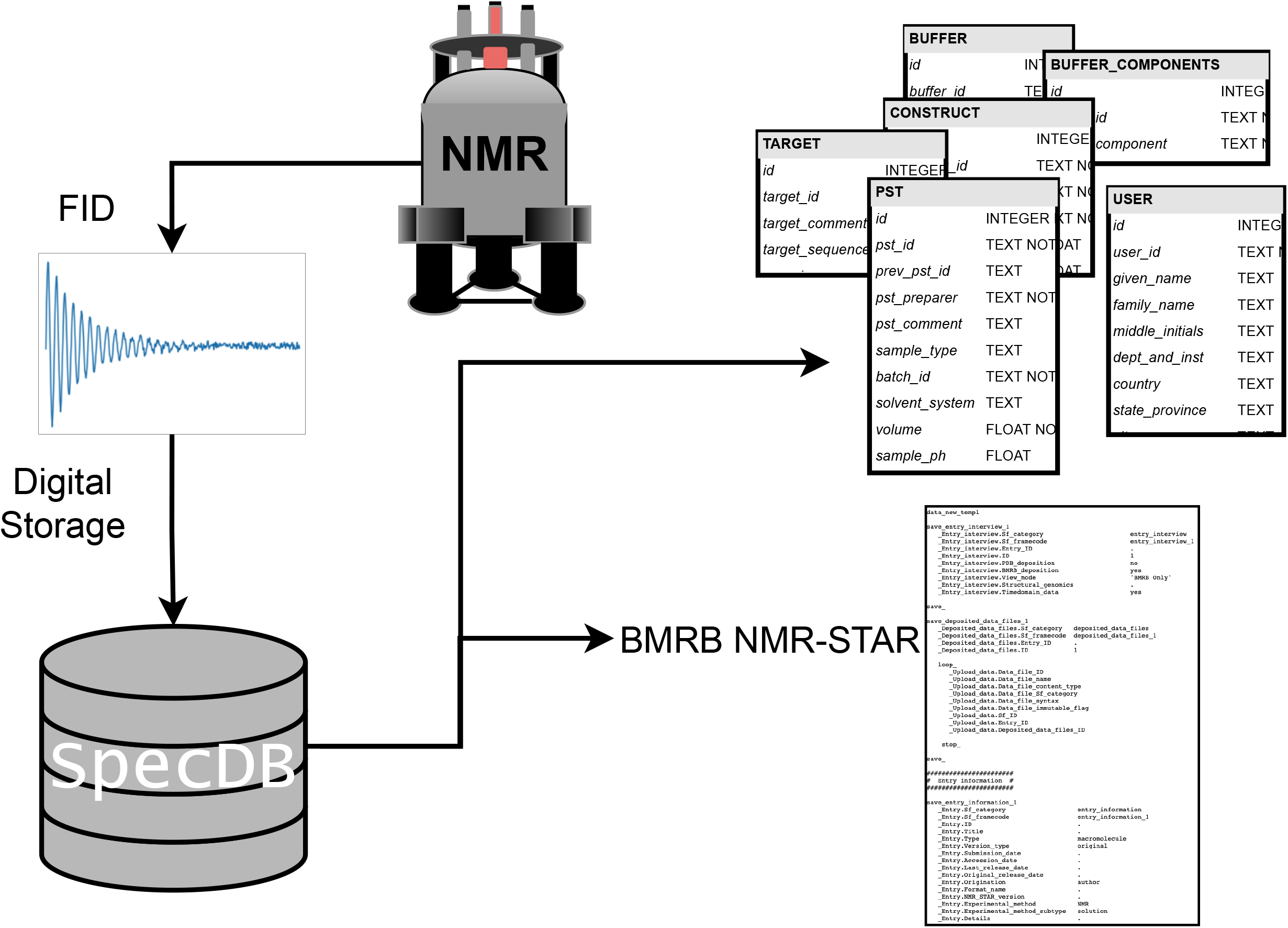

## Introduction

The success of machine learning in biology over the past 10 years, particularly deep learning in the field of protein structure prediction^1–4^, is leading many communities in the biological and medical sciences to reevaluate their data ecosystems^5^. Data is the key resource to train and deploy sophisticated machine learning models^6^, and the degree to which well-organized data is available can spur innovation across biology, chemistry, and medicine. Modern protein Nuclear Magnetic Resonance (NMR) spectroscopy laboratories also require an easy-to-use, lightweight data management system for managing NMR time domain data, archiving them locally, and eventually moving these data into public repositories. The goal of this study is to advance the data infrastructure and practices for the scientific community to provide tools and protocols for archiving NMR time domain data for future data mining and machine learning.

NMR spectroscopy has a rich history of developing and applying machine learning at all stages in the experimental pipeline^7,8^. High impact examples include predicting protein torsion angles^9^, chemical shift prediction^10^, NMR spectral peak picking^11,12^, and reconstruction of non-uniformly sampled free induction decays (FIDs)^13–15^. Additionally, there have been efforts to organize NMR data into datasets suitable for machine learning, like the RefDB dataset with re-referenced chemical shifts^16^. Designing deep neural network architectures and applications of existing deep learning methods to tasks across the NMR data analysis and structure determination pipeline is an active area of research. To further develop and engineer sophisticated machine learning methods for NMR data analysis requires an accessible data infrastructure to collect more and richer datasets.

The free induction decay collected from the NMR spectrometer is the raw data that all proceeding steps in an NMR experimental pipelines rely and build on^17^. The terms *time domain data* and *free induction decay* (FID) data are used interchangeably in the community for these raw data. The prospect of automatic analysis of FIDs to produce NMR resonance assignments, dynamic information, or even molecular structures, is a long-standing goal and challenge in the NMR spectroscopy field. A large set of curated time domain datasets is a critical first step to support such applications.

Biomolecular NMR time domain datasets are archived in the **<u>B</u>**iological **<u>M</u>**agnetic **<u>R</u>**esonance **<u>B</u>**ank (BMRB)^18^. However, only a small percentage of BMRB entries have associated time domain data. One way to address this data gap is to provide a simple tool to allow organization and archiving of FID data, and associated metadata, soon after they are generated at the NMR spectrometer, and to provide a simple process for moving these data into the BMRB. In this way, a data resource of FIDs will grow in time.

Our approach to addressing the challenges in archiving and distributing raw NMR time domain data is a data management tool called **<u>Spec</u>**tral **<u>D</u>**ata**<u>B</u>**ase. SpecDB is a simple data management system that individual NMR research groups can install and use to create their own archive of organized experimental NMR data. SpecDB also provides capabilities to share all, or selected sets, of these data between research groups, and to transfer these data to the BMRB. Furthermore, SpecDB should be easily maintained by any spectroscopist or NMR spectroscopy research group, without much relational database knowledge.

One important goal recommended by the wwPDB NMR Validation Task Force is to foster community practices of consistently depositing time domain NMR data into the BMRB^19^. The need for such large-scale efforts in preserving and disseminating FID data is greatly appreciated in the NMR and structural biology communities^20,21^. However, unless these time domain data are stored in an organized manner, together with appropriate metadata describing the sample and data collection parameters, deposition and retrieval of the underlying FID data for biomolecular NMR studies and machine learning is difficult and time consuming, and is often not even attempted. SpecDB provides a platform for organizing and storing NMR time domain data, together with metadata describing associated data collection parameters and samples, in a form suitable for future data mining and machine learning. SpecDB also addresses important issues of data reproducibility and validation of research results. This software platform is a step forward in developing a data infrastructure for learning on NMR time domain data, as well as promoting practices of regular deposition of FID data to the BMRB.

SpecDB is related to **<u>L</u>**aboratory **<u>I</u>**nformation **<u>M</u>**anagement **<u>S</u>**ystems, or LIMSs. There are, and have been, many LIMS developed by the NMR community, and across the chemical and biological disciplines. One successful LIMS is the **<u>S</u>**tructural **<u>P</u>**roteomics in the **<u>N</u>**orth**<u>E</u>**ast (SPINE) database^22,23^, built to support the protein sample production and structure determination efforts of the **<u>N</u>**orth**<u>E</u>**ast **<u>S</u>**tructural **<u>G</u>**enomics (NESG) Consortium (https://nesg.org/). The SPINE MySQL relational database tracks the progress of protein targets and projects through specific pipelines for protein sample production, characterization, and structure determination by NMR and X-ray crystallography. SPINE is associated with the OracleSQL relational database SPINS, **<u>S</u>**tandardized **<u>P</u>**rote**<u>I</u>**n **<u>N</u>**MR **<u>S</u>**torage^24,25^, the goal of which was to archive each step and associated data necessary to completely reproduce a specific protein NMR data analysis pipeline. Other successful LIMSs and/or software suites providing some of these same capabilities include ProteinTracker^26^, Sesame^27^, PiMS^28^, NMRFAM-SPARKY^29^, NMRbox^30^, and CCPN^31^ to name a few. SPINE and SPINS are specialized to support the pipeline and infrastructure of a specific pipeline of a large-scale structural genomics project, and are not sufficiently general, light-weight, and portable to support the broader needs for data archiving across the biomolecular NMR community. However, they serve as motivations and guides for the design of

SpecDB, which aims to address the specific data management problem of archiving NMR FID data and associated metadata by a small research group, needed to archive these FID data in the BMRB. SpecDB, an FID database suitable for use by a single laboratory or a biomolecular NMR facility, was developed with five principal features. (i) The raw time domain data (FID) is the centrally tracked entity. (ii) Experimentalists can also archive metadata items needed to describe the FID data through text forms. (iii) The system supports interchange between database items in SpecDB to database items tracked by the BMRB, to allow for BMRB deposition. (iv) The database is searchable with structured queries. (v) Query outputs can write FID and associated metadata from the SpecDB database into a folder-based hierarchy, to allow users to interact with the FIDs and sample information in a filesystem format.

In this paper we discuss implementation and system requirements for the SpecDB software, the relational schema in SpecDB, the overall workflow for archiving NMR FIDs using SpecDB, and some useful query tools. The software is freely available for implementation by any laboratory on Linux computer systems through the following GitHub code repository: https://github.rpi.edu/RPIBioinformatics/SpecDB.

## Methods

SpecDB is a software platform that can archive minimal sample and experimental descriptions of FID data obtained from an NMR spectrometer. The NMR spectroscopist can provide the appropriate information about the sample and NMR experiment in files or in text based forms. The information is then funneled into a relational database. With relational databases, data items are stored in tables, or spreadsheets, yet there are multiple tables where columns in one table connect or relate to columns in different tables. The relationships, or connections between columns in a SQL table is the relational aspect we are referring to. SpecDB is a database that NMR research groups can construct locally on their laboratory Unix or Linux computer systems. The SpecDB software has two overarching components: (i) the relational database that describes an NMR experiment data collection process and associated FID data implemented in SQLite, (ii) the Python software package of SpecDB that manages the insertion and querying of data from the database.

There are three key computational characteristics in SpecDB. First, the SpecDB schema and database is built using SQLite, a light-weight and fast implementation of SQL. SQLite powers many websites and scientific applications, and is an important industry standard in IT and data science. With SQLite, the entire relational database is a single file, which makes managing database read/write/query permissions equivalent to managing file permissions in a file system. Sharing within group(s) can be easily set up with group permissions. Second, the SpecDB code base is developed in the Python language, which is one of the most widely used programming languages, particularly in the data science and bioinformatics field. Third, SpecDB utilizes the **<u>J</u>**ava**<u>S</u>**cript **<u>O</u>**bject **<u>N</u>**otation (JSON) text interchange format^32^ to store key NMR experimental metadata items that describe an NMR experiment and FID data. JSON files are human readable, allowing investigators to easily work with them, and to update them interactively. Using JSON forms provides a general solution for representing metadata for biomolecular samples and NMR experiments. Various form filling tools can be developed and implemented in the future to produce the JSON files needed for SpecDB. In our current implementation of SpecDB, we use user-edited Google Sheets to create these JSON files. Fourth, SpecDB is developed for Linux operating systems as is common and standard in bioinformatics and structural biology.

## Results

Figure 1 illustrates the data ecosystem for protein NMR and the challenges with archiving and organizing the raw time domain data. NMR research groups typically use laboratory or institute NMR facilities. Within each NMR research group are individual investigators working collaboratively on diverse molecular systems and questions. It is often the case that the storage and organization of the raw time domain data is left to the individual scientist who collected the data, and this leads to many different practices and conventions, even within a single research group, for storing FIDs and the essential metadata that describe the experiment.

**Fig. 1:**
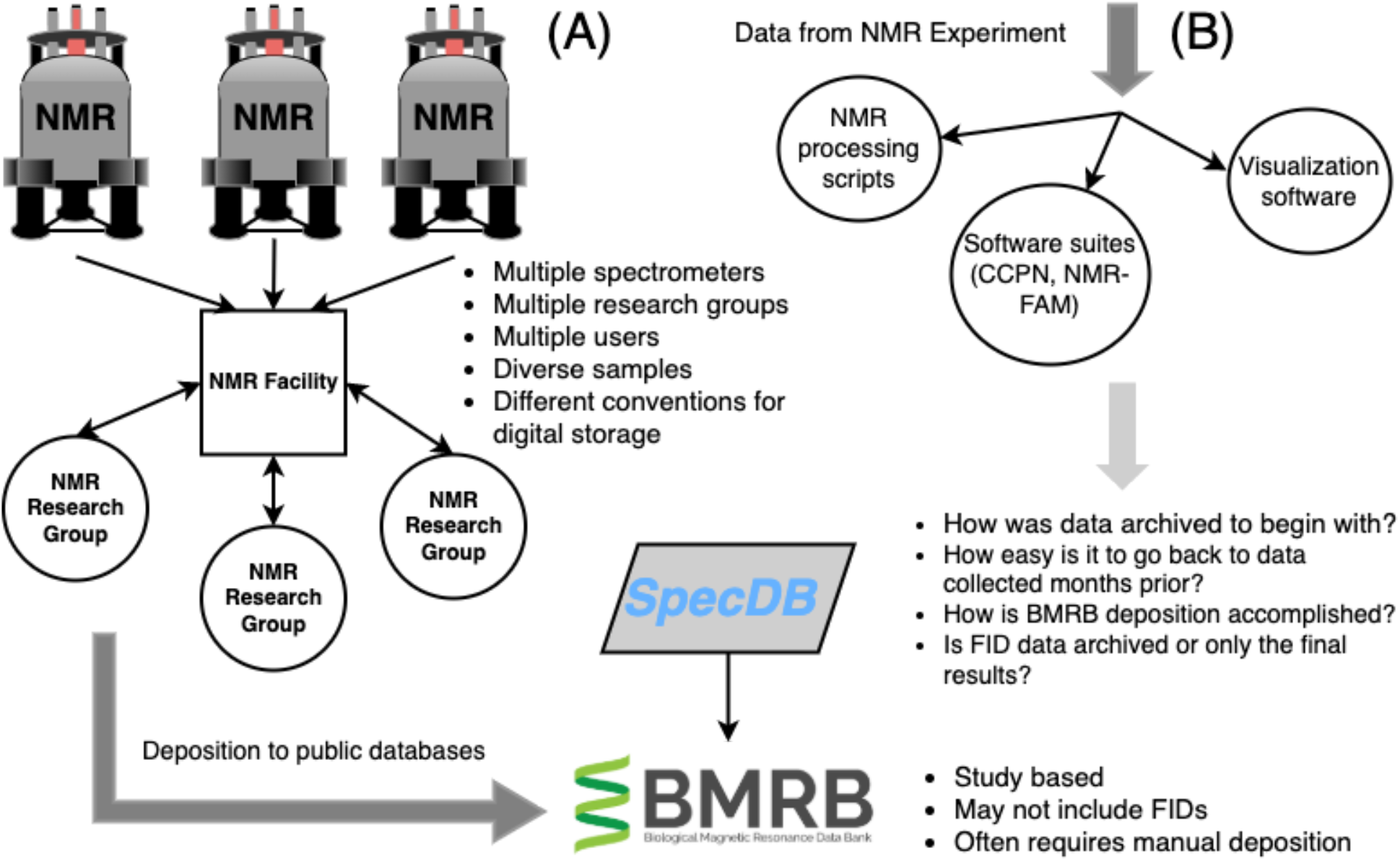
Data ecosystem for biomolecular NMR. (A) In general, biomolecular NMR research groups make use of shared NMR facilities where many NMR spectrometers are maintained and scheduled to specific users in specific research groups. After data is collected, a study is typically published and the experimental data to support the study is uploaded to the PDB and BMRB databases. The BMRB deposition is based on a specific study, and depositors are not required to submit time domain data. (B) Time domain data from NMR experiment is typically funneled into a processing phase of the data analysis, using specialized NMR processing tools, visualization tools, or other software suites for analysis and visualization, such as NMRPipe, SPARKY, CCPN, and NMR-FAM software suite. SpecDB provides solutions for some key questions including: how is the raw time domain data collected and stored?; how easy is it to find FIDs from a particular study or data range?; how do I retrieve the FID together with metadata and organize it for a BMRB deposition?

On commercial NMR spectrometer systems, the NMR FID data is included in a data collection directory that also includes many details of data collection, including the actual NMR pulse sequence code, spectrometer shim parameters, specific data collection parameters, pulse sequence waveforms, etc. In SpecDB, the “FID data” that is stored refers to this entire data collection directory. While most of the data items in these parameter files do not have specific representations in the SpecDB schema, they are still stored in the SpecDB database as a compressed directory. This allows for future development of the SpecDB schema to include specific data collection parameters, such as NOESY mixing time values or pulse widths, that are stored in these data collection directories. Hence, the initial focus of the SpecDB schema is to provide a platform for archiving these data, along with metadata about the NMR sample and other data collection parameters that are not included in these FID data directories.

### Process of Developing the SpecDB Schema

The SpecDB schema developed to describe FID data (actually, the FID data directory), NMR sample, and associated metadata is designed to be compatible with both the SPINE database schema^22,23^, and with the NMR-STAR data ontology^33^ used for archiving NMR data in the BMRB. Tables in the SPINE schema provide detailed information about protein samples, including information about the protein itself, the protein families it has been classified into, information about homologous proteins, disorder predictions, details of cloning, expression, crystallization data, and progress in structural characterization by NMR and X-ray crystallography. Most of these details are not required for SpecDB. We assessed each data item and data table in SPINE to identify a condensed subset that could minimally and routinely describe an NMR FID data collection experiment and the corresponding NMR sample. In addition, by inspecting numerous representative NMR-STAR files from the BMRB, and in consultation with the BMRB developers, we identified additional data items that need to be provided for deposition an FID dataset into the BMRB, and hence need to be tracked through the SpecDB database. SpecDB thus provides direct translation from SpecDB data items to NMR-STAR tags. By using both SPINE and NMR-STAR, we were able to arrive at a minimal SQL schema to describe an NMR FID dataset sufficiently well to ensure its reproducibility, and to convey it into the BMRB.

The schema of SpecDB can be viewed as having two main parts, the database tables that describe the NMR sample, and the database tables that describe the FID data. Figure 2 depicts this two-wing structure of the SpecDB schema. Making a simple schema that is general enough for a wide range of applications is a significant challenge. Hence, the SpecDB schema is designed to be flexible enough to provide for significant modifications needed to support specific data pipelines and query requirements. Some examples of information not included in the SpecDB schema include details about the DNA cloning protocols used for making protein constructs, details of biomolecule purification procedures, detailed information about fermentation and expression, and bioinformatics, evolutionary, and gene-family metadata about the biomolecular target. These are not essential for the process of archiving the FID data and depositing it into the BMRB. However, the schema provides the flexibility for expansion to handle these additional data items in the future, which can be guided by, for example, the SPINE schema which includes many of these additional sample preparation details.

**Fig. 2:**
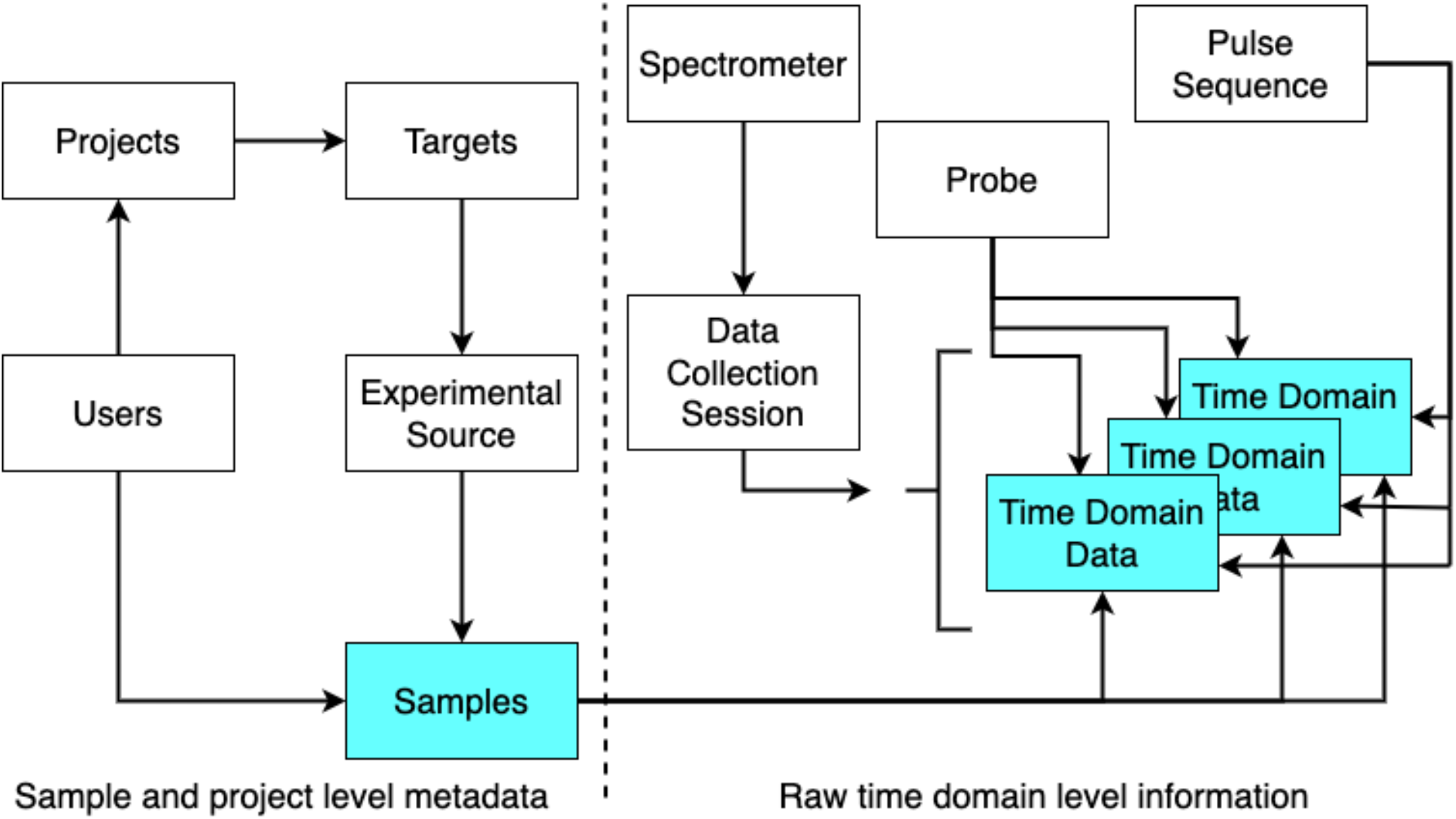
The two wings of the SpecDB schema. The SQL schema for SpecDB can be considered in two parts, or wings. First is the description of the experimental Sample used for NMR data collection (left side). Users define Projects, Targets, Experimental Sources, and Samples. A Sample is part of a Project, defined by the group using SpecDB. Within Projects are Targets, biomolecules that are the subjects of the Project study. Experimental Sources describe aspects of the production of the Target. Samples (PSTs) are the actual samples that are analyzed at the spectrometer. The second wing of SpecDB relates information about the FID data (right side). SpecDB collects information about the Spectrometer, Probe, and Pulse Sequence used for collecting a specific FID. On some spectrometry systems, including Bruker systems, FIDs are collected in a “Session”, which is a series of related NMR experiments. This session hierarchy is preserved in the schema of SpecDB. The right-hand side of the figure indicates the many-to-one relationship between Sessions and FIDs, as there are multiple time domain datasets associated with the Sessions table.

### SpecDB Tables that Provide Sample Information

The main table in SpecDB that describes the NMR sample is the **<u>P</u>**rotein **<u>S</u>**ample **<u>T</u>**ube (PST) table. The Protein Sample Tube refers to a physical tube holding a sample (which may be protein, nucleic acid, or other biomolecule or non-biological chemical). It includes sample tubes used in preparing a sample (e.g. Eppendorf tubes), or the actual NMR tube inserted into the NMR spectrometer. The set of relational tables that specify the sample and project are summarized in Figure 3. Each Protein Sample Tube is assigned a unique text identifier by the user called the *pst_id*. The *pst_id* is assigned by the user/research group. This identifier must be unique in the database. However, the actual id is determined by a lab-specific naming convention. This naming convention system is discussed below.

**Fig. 3:**
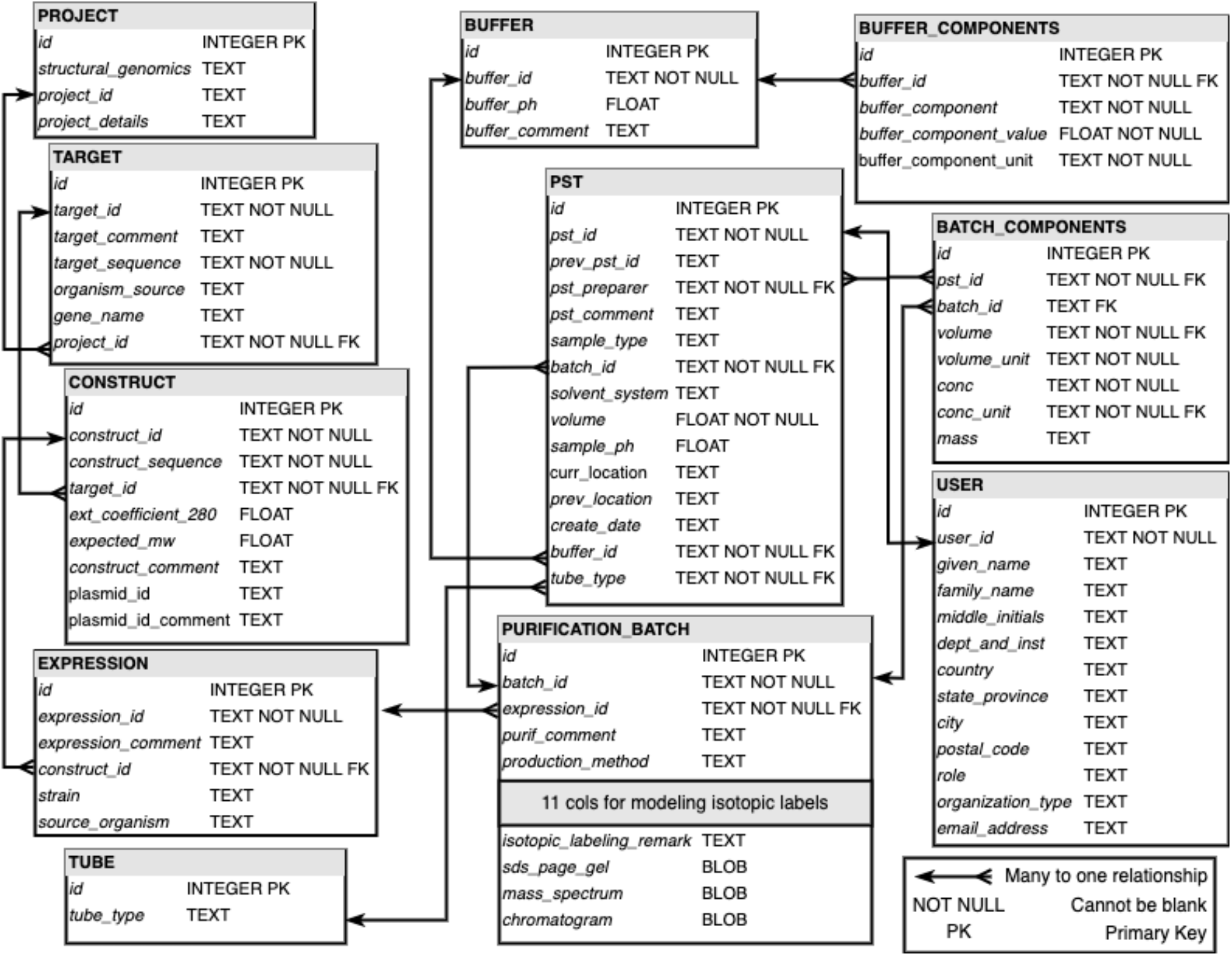
Relational diagram for SpecDB tables. A view of the tables that describe an NMR sample and project. A hierarchy of meta-data information is depicted in the nested relationships between PROJECT, TARGET, CONSTRUCT, EXPRESSION, PURIFICATION_BATCH, and PST. Across the entire SpecDB schema there are 17 tables, 12 of which are displayed above for the description and modeling of NMR samples. Some data items (e.g. isotope-enrichment tags, shown in Supplementary Table S1) are excluded for clarity. The connectors between tables indicate the relationships between tables. All the connectors in the diagram indicate a specific type of relationship, many-to-one relationships.

A key feature of the sample specific tables presented in Figure 3 is the nested nature of these tables. A sample description starts with the PROJECT table. Samples are part of a project, or cohesive study. The data items in the PROJECT table describe the research project, and provide a simple unique name for the project, the *project_id*. The hierarchical flow of information for describing a sample follows PROJECT, TARGET, CONSTRUCT, EXPRESSION, PURIFICATION_BATCH, BATCH_COMPONENTS, and PST. Multiple samples, each prepared as a “purification batch”, may be combined in a single PST to form complexes, as defined by the BATCH_COMPONENTS table. As a consequence of this hierarchy, every purification batch is associated with an expression experiment (also called a fermentation) run, every expression experiment is associated with a construct, every construct is associated with a target, and every target is part of a project. This nested hierarchy reflects the SPINE data schema, and in the future will allow for archiving NMR spectra from the NESG SPINE and SPINS databases into SpecDB for public distribution.

Inspection of the tables in Figure 3 illustrates that nearly every table has a text based identifier that is unique across the respective table. For example, the PST table has a *pst_id*, which provides a unique name for each protein (or nucleic acid) sample tube. SpecDB does not impose a specific convention or nomenclature on the data record identifiers (*project_id, target_id, pst_id*, etc), except that each data record must have a unique identifier. The naming convention for these unique identifiers should follow a convention set by the research group. For example, assignment of the unique textual identifier for a Protein Sample Tube, *pst_id*, may be chosen by the user who prepared the sample tube. There is no internal SpecDB mechanism to generate identifiers other than persevering their uniqueness within their respective table. However, SpecDB checks identifiers at data input to prevent using an ID already in the database (unless a user specifies with a flag the need to update the associated record).

In the SPINE database, record id’s follow a convention based on the *project_id*; e.g. HR for “human protein project at Rutgers”. At each subsequent level in the organization hierarchy (targets, constructs, expressions, purification batches, and PSTs), there is a new delimiter that is added to the ID to make the ID unique and convey some information about the sample. Accordingly, *project_id* name HR defines the *target_id*’s; e.g. HR001A (the first domain of 1st protein HR001, in the project HR), which then defines the naming of *construct_id’s, expression_id’s, purification_batch_id’s, and pst_id’s*. In this example, the *purification_batch_id* HR001A.200_345_NTag.NiNTA.004 is the 4th batch of a construct of target HR001A that comprises residues 200 - 245 with an N-terminal hexaHis tag purified by NiNTA affinity purification. The corresponding NMR sample tube *pst-id*’s are assigned abbreviated names based on the *target_id* (e.g. HR001A.001, HR001A.002, etc.), which fit better on NMR tube labels. It should be noted that this naming convention is convenient, but does not replace accessing the corresponding PST record (and the associated hierarchy of records) to get complete and accurate information about the sample. Users of SpecDB may adopt this convention, or develop their own unique naming system for record ids.

Inspection of the schema for the PST table (see Figure 3) highlights the relational nature of SpecDB. The PST table links to other tables that describe the sample. For example, the user who generated the Protein Sample Tube is recorded in the PST table using the *pst_preparer* column. The value in the *pst_preparer* indicates the user who created the PST. In order to know the many attributes that describe a user, the value provided in the *pst_preparer* column is a key that links back to the USER table where the remaining items to describe the user are stored, rather than creating many columns within the PST table to record the user’s first name, last name, email address, etc. All the user’s information is stored in the USER table, and is linked to the necessary rows in the PST table through the *user_id* key. There are four tables that all connect to the PST table, as illustrated in Figure 3 through the barbed connectors. The barbed connectors indicate that the relationship between the two tables being connected is a many-to-one relationship. For instance, many sequence constructs can be made for a single protein target.

Next we will describe each table presented in Figure 3 and the role each plays in describing an NMR sample, following the hierarchy discussed above. A target is generally a biomolecule (protein, nucleic acid, polysaccharide, etc), although non-biological molecule samples can also be described with this schema. The biomolecule may come from a natural source, or be artificially designed or synthesized. For natural proteins, the protein is defined by the Uniprot^34^ protein sequence of the full-length protein. A unique *target_id* is defined for each target, and linked back to the corresponding PROJECT table. Following the TARGET table is the CONSTRUCT table. It is often the case that the biomolecules being studied with NMR have an amended primary sequence, for purification reasons (e.g., a purification tag), resulting from mutations introduced for functional studies, due to truncations to suppress aggregation, or for other reasons. Hence, the construct sequence studied by NMR is generally different from the target sequence. A construct is assigned a *construct_id* and a link to the *target_id* from which it was made. Associated with each construct are one or more expression (or fermentation) experiments. The EXPRESSIONS table, designated by a unique *expression_id* provides metadata on how the expression of the construct in a particular bacterial strain or other organism was accomplished.

Following expressions are protein purification batches, described in the PURIFICATION_ BATCH table. This table also provides a *sample_sequence*, which may be different from the construct_sequence if purification tags are removed in the process of purification. Here, SpecDB also allows users to store the absorbance extinction coefficient (e.g., at 280 nm for proteins) expected for the purified sample_sequence, which can be estimated relatively accurately from the protein or nucleic acid sequence^35,36^, and the expected molecular weight. If the construct is isotope-enriched, this needs to be accounted for when retrieving the expected molecular weight from the sequence.

Within the PURIFICATION_BATCHES table is a recording of the isotopic-labeling actually achieved for the biomolecule. The isotope-labeling may be that expected based on the isotope-enrichment strategy used, or that determined by experimental data such as NMR or mass spectrometry. Not all the isotopic labeling schemes tracked in the SpecDB schema are listed Figure 3; additional schemes are listed and described in Table S2. Currently, eleven common types of isotope labeling can be tracked in the PURIFICATION_BATCH table and more can be added as needed in future versions. The *isotope_labeling_remark* is a free-text field that allows the user to record labeling methods not captured by the isotope-enrichment strategies currently supported for the PURIFICATION_BATCH table. A PST may come from a single purification batch, or (in the case of complexes) multiple purification batches. The one or more batches combine to form a PST are tracked by the BATCH_COMPONENTS table.

The PST table also provides a description of the protein sample tube itself using a controlled vocabulary of common sample and NMR tubes, including conventional NMR tubes and Shegemi NMR of various diameters (i.e. 1-mm, 1.7-mm, 3-mm, 4-mm, 5-mm, 8-mm, 10-mm). In the case of solid-state NMR, a PST tube can be a rotor of various sizes. The PST also tracks the actual sample pH (or the expected pH based on the buffer used), who prepared the sample tube, and the physical location of the protein sample tube, the solvent, the buffer, as well as the the sample volume and concentration of the target molecule(s) in the sample tube.

Associated with the PST table is the BUFFER table. The BUFFER table records all the buffers used in the database, and each buffer is provided a *buffer_id*. A *buffer_ph* is recorded, which may be different from actual *sample_ph* recorded in the PST table. In order to describe the contents of a buffer, SpecDB also has a BUFFER_COMPONENTS table. Each row of the BUFFER_COMPONENTS table is a different component used to make buffers, where the buffer is associated with this component through the *buffer_id*. A buffer component requires three items to complete its description: the name of the component, the concentration of that component, and the unit of concentration. Buffers can be very complex, and having a simple table structure to record all the buffer components of a particular buffer may be tedious in the short term, but highly valuable due to accuracy in archiving the sample, reproducibility, and for future data mining.

The last table to highlight in Figure 3 is the USER table. Here, the investigators in the research group are recorded, their names, emails, department and institution, etc. The USER table is important for many reasons. In particular it is helpful for trouble-shooting a project when it is known who made a particular sample, or recorded a specific spectrum. User information is also required for creating a BMRB deposition, and for documenting credit for publication.

Elements of the SpecDB schema not illustrated in Figure 3 are the controlled vocabularies on the SpecDB data items, or the text strings or values allowed to be inserted into the database. Not every data item has a controlled vocabulary, but several require controlled vocabulary to ensure consistency in what users input as information across the schema. Table 1 presents a representative sample of the data items that have a controlled vocabulary in the SpecDB schema. As an example, items such as *volume_unit* cannot take any text string, there are only certain text strings (i.e., units of volume) allowable to be used for the *volume_unit* value. This helps maintain consistency in the database.

**Table 1:** Controlled vocabularies across the SpecDB relational schema. These are representative examples of the controlled vocabularies used by SpecDB. Indicated are the SpecDB data types where a controlled vocabulary exists, a description of the data modeled for each data item, and the exact expression that controls the allowable values for each SpecDB column. Not every column with a controlled vocabulary is presented in this table. The allowable tube type names are listed in in Table S1, and controlled vocabularies for different isotopic labeling is described in Table S2.

| SpecDB data type | Description | Controlled Vocabulary Description |
| --- | --- | --- |
| *structural_genomics* | Optional for BMRB deposition. Indicate if project is part of a structural genomics effort. | ( 'yes' OR 'no' ) |
| *iso_13c_enrichment* | Describe the carbon-13 isotopic enrichment. | Must have substring ( " % 13C " ) |
| *buffer_component_unit* | Record the unit the buffer component exists in the buffer. | ( 'mM' OR '% (v/v)' OR 'mg/ml' ) |
| *sample_type* | Record the type of PST sample. | ( 'solution' OR 'solid' ) |
| *volume_unit* | Unit of volume for protein component of PST. | ( 'nL' OR 'μL' OR 'mL' OR 'L' ) |
| *conc_unit* | Concentration unit for protein component of PST | ( 'μg/mL' OR 'mg/mL' OR 'nM' OR 'μM' OR 'mM' ) |
| *nus* | Indicate if non-linear sampling was employed | ( 'yes' OR 'no' ) |

### SpecDB SQL Tables to Archive FIDs

The second wing to the SpecDB schema are the tables that describe FID data sets (Figure 4). An FID is recorded on a spectrometer, so there is a SPECTROMETER table that records the names of the spectrometer used, the spectrometer model, and the field strength.

**Fig. 4:**
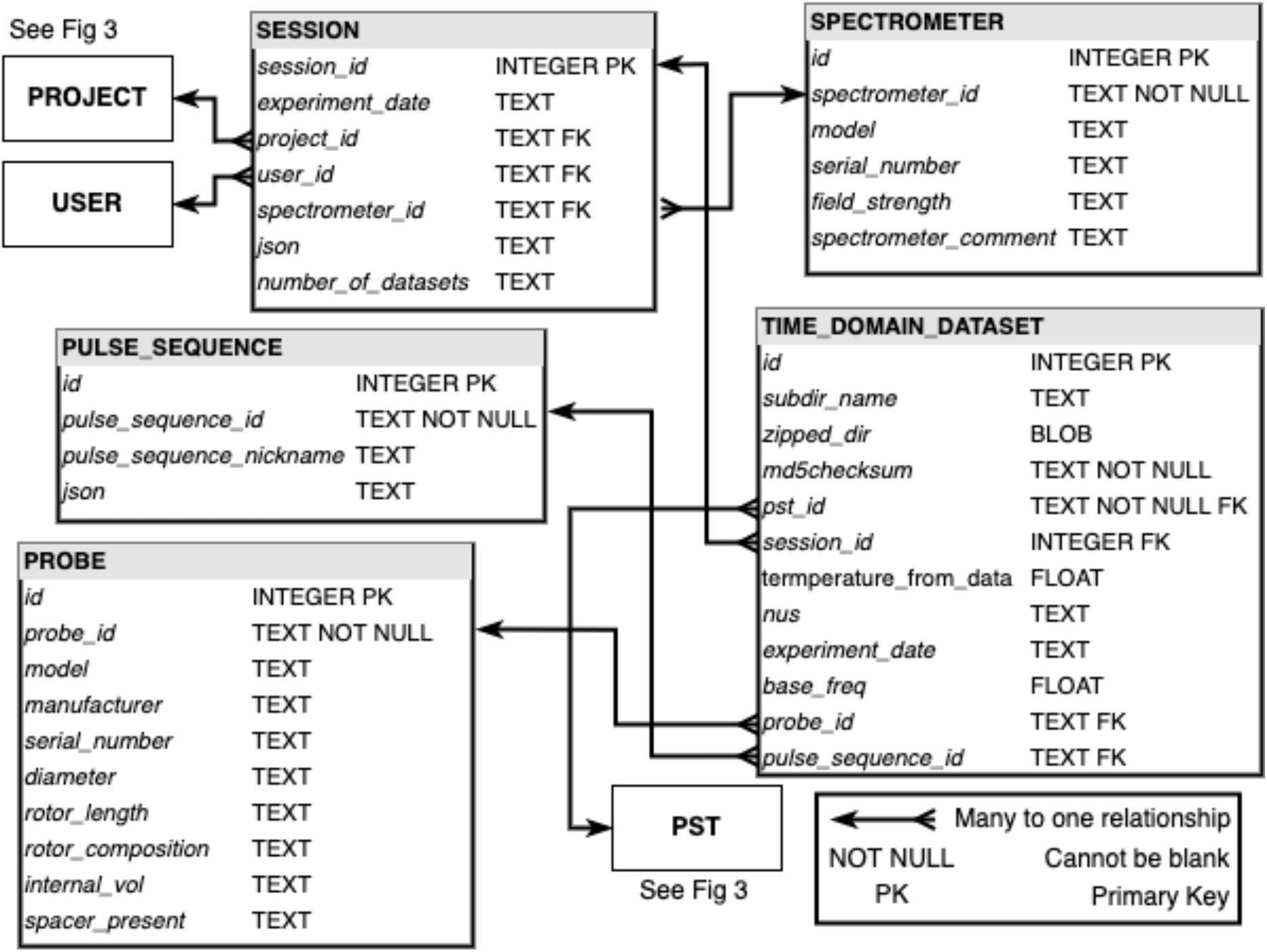
Relational diagram for SpecDB tables that describe NMR FID data. The relationship diagram depicted in this figure are for the tables in the SpecDB schema that describe NMR experiments and the data collected at the NMR spectrometer. Inside the diagram are callbacks to the PROJECT, USER, and PST tables in Fig. 3. The complete FID subdirectories are stored as **<u>B</u>**inary **<u>L</u>**arge **<u>Ob</u>**ject**<u>s</u>** (BLOBS) in the *zipped_dir* column of the Time_Domain_Datasets table, allowing other auxiliary files such as acquisition/acquisition status files needed to reproduce the experiment to be archived along with the time domain NMR data.

The next level of this hierarchy is SESSIONS, which are sets of FIDs collected together in a data collection session. A session could be a single FID data set (as for example data collected on Varian spectrometers using VNMR software), or a directory containing subdirectories with a single FID in each (as is the case on Bruker spectrometer systems). The concept of a session stems from the management of FIDs on Bruker spectrometers using TopSpin software. Here, the NMR spectroscopist may queue up several pulse sequences to be run in succession at the spectrometer. The FIDs from these pulse sequences are placed into different subdirectories of a session directory. This subdirectory structure is reflected in the SpecDB SESSIONS table. In the SpecDB SESSIONS table, the spectrometer that is being used is recorded, the data collection dates, the number of FIDs to be collected in the session, the project associated with the session, along with the user running the session. Internally to SpecDB, each session is given a *session_id*, which is simply a row integer counter, and is an item that the user does not set. The session directory contains the *specdb*.*json* JSON file, with metadata provided by the user, as well as the sub-directories with the recorded FID data and spectrum-specific acquisition parameters. In this way all of the metadata describing all of the FID data collected in the session, including information about the user, sample, and other aspects of the data collection are all stored together in the JSON text file at the session directory level.

The SESSIONS table is a useful LIMS concept for NMR data management. Sessions reflect the fact that some FIDs are related to other FIDs. The SESSIONS table also highlights that SpecDB is more than a database of FIDs, it describes the samples the FIDs are recorded from, and also maintains information about relationships between FIDs.

Ultimately, the recorded FIDs (i.e. FID data directories) themselves are stored in the *zipped_dir* column of the TIME_DOMAIN_DATASETS table. The *zipped_dir* data item is a compressed data directory containing the FID and the associated vendor-specific data collection metadata. The *zipped_dir* is stored in the database as a **<u>B</u>**inary **<u>L</u>**arge **<u>Ob</u>**ject (BLOB). To ensure that every stored FID in SpecDB is unique, we perform a MD5 hashing function on the raw FID data file to be inserted to SpecDB, and the hashed string is stored in the TIME_DOMAIN_DATASETS table in the *md5checksum* data item.

Also included in the TIME_DOMAIN_DATASETS table is the *probe_id*, which links back to the PROBES table that describes the NMR probe used in the collection of the FID. The probe information is stored at the level of the FID instead of the session in the SpecDB schema because it is possible that within the same session, users may switch out probes for different applications. The probe information collected in the PROBES table displayed in Figure 4 contains items that relate to probes for both solution and solid-state NMR.

The TIME_DOMAIN_DATASETS table also includes a name of the pulse sequence used to collect the corresponding FID, and the *pst_id* for the sample being analyzed. The pulse sequence name is a nickname (e.g. 2D NOESY) used for queries or BMRB deposition; the actual pulse sequence miniprogram is included in the FID data directory associated with the TIME_DOMAIN_DATASETS table. Like the probe information described above, the *pst_id* is modeled at the level of the FID instead of session because spectroscopists might change the sample, or use different samples over the course of a session. For instance, spectroscopists might perform a pH titration by adjusting the pH in the sample tube and recording a new FID on the adjusted sample tube. In this case, each time the pH is changed results in a new *pst_id*. The spectroscopist records the path of sample changes using the *prev_pst_id* data item in the PST table, allowing protein sample tubes to inherit and trace information from each other.

There are two remaining tables that are not represented in Figures 3 and 4 and do not necessarily fit the two-wing structure used to describe the SpecDB schema. The first is the STAR_CONVERSION table. This table is not intended to be modified by users as it contains a translation between the SpecDB data items to NMR-STAR tags. This helps the SpecDB applications for writing database contents into NMR-STAR formats. The second table, discussed below, is the SUMMARY table for queries, which is a subset of commonly searched data items in SpecDB in one flat table.

### SpecDB Workflow

The intended workflow with SpecDB is illustrated in Figure 5, which depicts an NMR spectrometer with the associated computer workstation where the FID is initially recorded. Typically, these FIDs cannot be stored indefinitely at these NMR workstations, and are moved to a laboratory server where SpecDB is installed, using *rsync* or other mirroring operation. SpecDB is run and queried on this laboratory server.

**Fig. 5:**
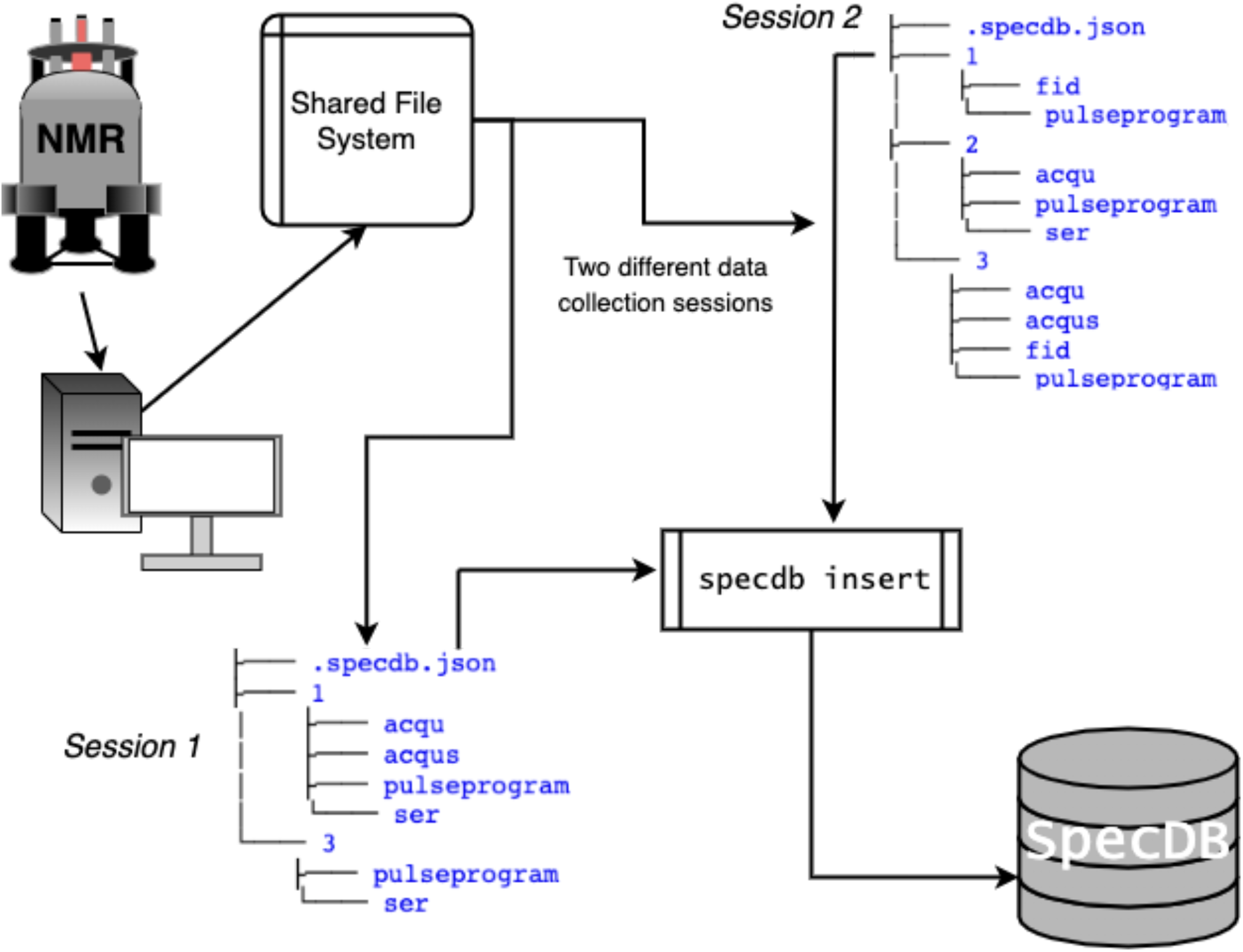
Movement of NMR time domain data from NMR spectrometer to SpecDB. FIDs are generated at the spectrometer and stored on the associated computer workstation. Typically the collected data from the NMR spectrometer workstation is then transferred, either through an *rsync*, a mirror, or manual copying to a different, more stable filesystem. The NMR spectroscopist will typically store the FIDs they collect in a directory somewhere in the shared file system. These directories are highlighted with the indicated *Session 1* and *Session 2* directory structures. From both *Session 1* and *2*, there are two or more sub directories with FID data (denoted here as fid for 1D NMR data and ser for multidimensional NMR data). At the top level of these sessions sits a *specdb*.*json* JSON file. The *specdb*.*json* describes the data collection session. Once the spectroscopist enters the required metadata information into the JSON file, the *specdb insert* command is used to insert the *specdb*.*json* file into the database.

The JSON file (*specdb*.*json*) is located in the main directory of a data collection session, and contains information about each FID data set in the subdirectories under that session. The structure of the JSON file defines which sample, pulse sequence, etc. is associated with each subdirectory FID data set (Figure 5). The JSON file may be edited either before data collection (e.g, entering sample data prior to data collection), at the spectrometer (e.g. designating which NMR experiment is being collected in each subdirectory), or after moving the data to the database server (e.g. completing metadata information prior to submitting the data to the database). Once the JSON file metadata is complete, the SpecDB command line tool can be run by the user to insert the FID data sets and metadata for the session into the SpecDB database.

### SpecDB Sub Commands

There are six subcommands in SpecDB: *create, backup, restore, insert, forms, summary*, and *query*. Table 2 lists each SpecDB subcommand, the arguments each takes, and an example command as potentially run at the command line. The SpecDB subcommands *create, backup*, and *restore* are designed to be used by a research group’s SpecDB manager. A new SpecDB database is created with the command *specdb create*. The location where the SpecDB SQLite database resides, and the backup SQLite database file are command line arguments to the *create* subcommand. Together, *specdb backup* and *specdb restore* perform the incremental backup operations for SpecDB. The subcommands *insert, forms, summary*, and *query* are intended to be routinely used by individual researchers. The SpecDB command-line tool allows users to interact with the database in a shell environment, but we are also developing graphical applications to make these commands more user-friendly.

**Table 2:**
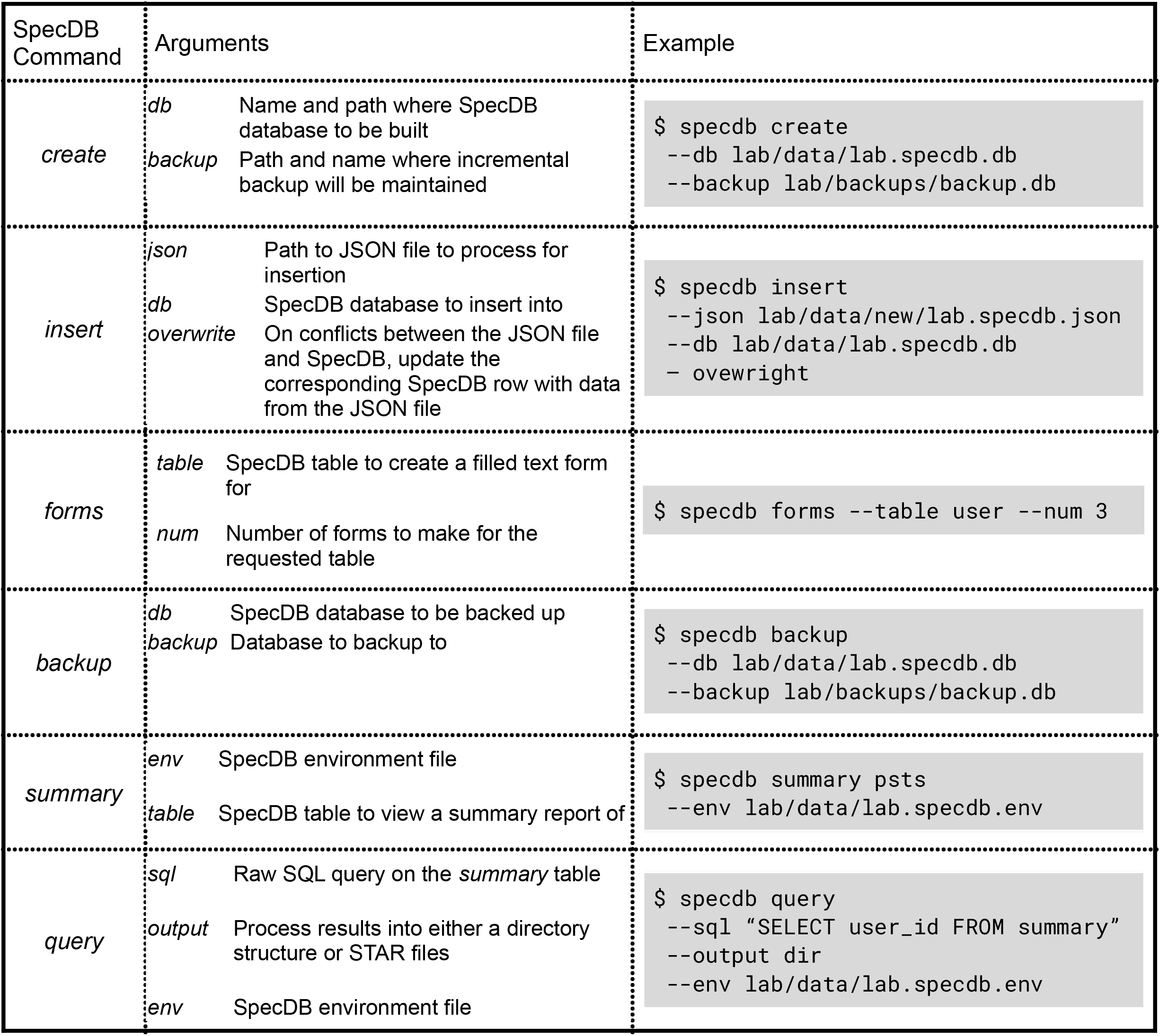
Description of SpecDB subcommands. The above table lays out all the commands within the SpecDB library that are used to manage NMR FID data in a filesystem and a SQLite database. The left column provides each sub command name. The middle column provides documentation on the command line arguments for each sub command. The right most column provides illustrative examples of how each SpecDB sub command could be executed in a general shell environment.

Once the user has completed a *specdb*.*json* file for their data collection session, these data are inserted into SpecDB database using *specdb insert*. Inserts that would override data already present are not allowed by default: SpecDB warns the user and forces the user to confirm if editing of previous values is intentional. s*pecdb forms* can be used to generate template JSON forms for any data item/table in the database, to provide a guide to assist users in creating a JSON file of metadata..

The *specdb summary* subcommand provides a summary of any table in the subject SpecDB database instance. Using *specdb summary* the contents of the requested table is printed in a formatted table. For example, *specdb summary users* will display a formatted table in the terminal session with table columns *user_id, given_name, last_name*, etc, and rows be the users that have been entered into the database. This allows users to review the data items and their values already inserted into the database, which can help users complete their *specdb*.*json* files and to assess inconsistencies.

Lastly, *specdb query* command allows users to perform queries against a SpecDB database and retrieve the subset of FIDs data sets that satisfy the query. These data are output in one of two formats, either a directory hierarchy of the data or NMR-STAR files for each FID. Building a SQLite database for NMR FIDs and sample information allows researchers to utilize the SQL language to extract data from the database using diverse and complex queries using SQL. The *specdb query* tool is designed to give researchers a way to make queries against a SpecDB database without using a sophisticated SQL query. With *specdb query*, users submit a SQL *SELECT* statement to be run against a SpecDB database. The *specdb query* tool will return all FIDs captured in the provided SQL *SELECT* statement. However, *specdb query* will only accept queries of data items listed in the SUMMARY table. SUMMARY is a SQL view of the SpecDB database, where columns from different tables are stitched together into a 2-dimensional table that is compatible with spreadsheets. More complex queries can be accomplished by connecting directly to the SpecDB SQLite database file. Table 3 lists out the exact terms incorporated into the SpecDB SUMMARY view, as well as examples of each data item.

**Table 3:** Schema description of SpecDB *Summary* View. This table presents the specific items tracked in *Summary* View in the SpecDB schema. Users can make structured queries against these columns and elect to have the query results be formatted into a directory structure or into NMR-STAR files. Left column indicates the names of the columns in the SpecDB *summary* view. Middle column is a description of each column in the *summary* view. Right column provides an example of the data types stored in each of the columns.

| Column Name | Description | Example data |
| --- | --- | --- |
| *id* | Row counter that assigns a unique integer to every FID collected | 12 |
| *experiment_date* | Data FID was collected | 2022-01-11 |
| *user_id* | The *user_id* of the user that collected the FID | KJF |
| *project_id* | Project_id for the project the FID is a part of | SPIKE Project |
| *structural_genomics* | Whether the FID is part of a structural genomics project | no |
| *temperature* | Temperature FID was collected at | 25 C |
| *buffer_id* | Buffer identifier that sample was in | NMR-Buffer-17 |
| *pst_id* | Protein Sample Tube identifier for the sample the FID was recorded of | SPIKE.2022 |
| *batch_id* | Identifier for purification batch | SPIKE.2022.b |
| *expression_id* | Identifier for expression run | SPIKE.2022.e |
| *construct_id* | Identifier for construct | SPIKE.102-450 |
| *target_id* | Identifier for target | SPIKE |
| *target_sequence* | Protein sequence of target | MGHSSSTVLAM... |
| *construct_sequence* | Protein sequence of construct | HHHHHHLEMGHSSSTV... |
| *pulse_sequence_id* | Name of pulse sequence | 1H-NOESY |
| *spectrometer_id* | Identifier of spectrometer FID was measured at | Hu800 |
| *field_strength* | Field strength of spectrometer | 800 MHz |
| *probe_id* | Identifier for the probe used in FID acquisition | Avance_2033478 |
| *tube_type* | Type of tube PST was in | 4-mm Shigemi tube |
| *nus* | Whether non-linear sampling was employed in FID acquisition | no |
| *zipped_dir* | Binary object of the zipped Bruker directory that contains the FID | (BINARY) |

Figure 6 illustrates a *specdb query* and shows condensed examples of the two output format types. NMR researchers are often expecting a directory structure when they are working with their data, so outputting query results as a directory hierarchy is a natural format option. One goal of SpecDB is to also generate FID data sets and metadata in NMR-STAR format by a query against the database. Using the STAR_CONVERSION table, every SpecDB data item can be translated to NMR-STAR save frames and tags. These NMR-STAR files can be used for deposition to the BMRB, or for sharing experiments between researchers and labs.

**Fig. 6:**
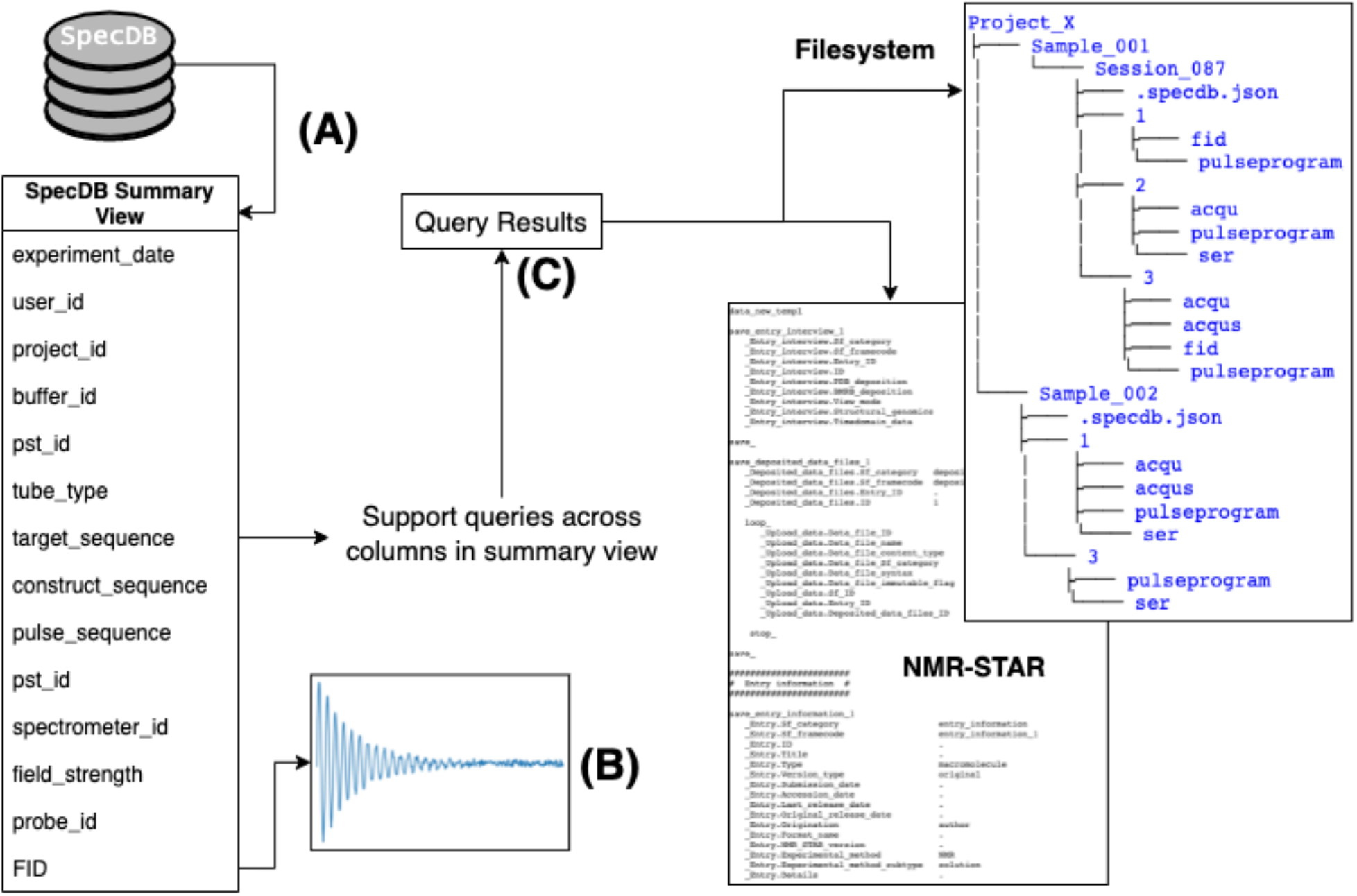
Overview for the SpecDB query system. (A) The list on the left of the figure above is a condensed version of the SpecDB Summary table; the complete list of columns supported in the Summary view is provided inTable 3 Supplementary Table X.. (B) A link to the raw binary data for each free induction decay (FID) is included in the Summary view. (C) SpecDB restricts SQL queries to data items in the Summary view. More complex queries can be handled directly through *sqlite3*. Queries generate FID data collection directories, formatted either in a filesystem folder hierarchy or as a set of NMR-STAR files.

## Discussion

SpecDB introduced in this study is a lightweight, flexible, robust LIMS for organizing and archiving NMR FID data generated in a small NMR research group or a large NMR facility center. In this first iteration of SpecDB, we had five goals: (i) Archive time domain FID data and key associated metadata, (ii) harvest user-supplied metadata that describes an FID experiment in human read-able JSON files, (iii) provide tools to allow queries of FID data sets in the database, (iv) allow records in SpecDB to be queried, organized, and formatted in NMR-STAR format for automatic deposition to the BMRB, (v) allow users to query, organize, and output SpecDB contents in a user-friendly hierarchical directory structure. All five goals are successfully implemented in version 1.0 of SpecDB. The SpecDB schema, based on the SPINE and NMR-Star schemas, was developed to describe an NMR sample and FID data set. Although focused on supporting descriptions of biomolecular (e.g. proteins and nucleic acids), the schema will also support non-biological NMR sample descriptions. SpecDB also includes command line tools that manage the insertion of new data into the SpecDB database, incremental backup of the database, and querying and retrieval of data from the database.

SpecDB falls under the general umbrella of a **<u>L</u>**aboratory **<u>I</u>**nformation **<u>M</u>**anagement **<u>S</u>**ystem (LIMS). There are several LIMS systems for NMR studies, and for many other domains of science. The diversity of LIMS systems is driven by the unique needs of a scientific discipline and community data standards. Dedicated LIMS have been developed for individual research groups that have specific workflows. The challenge with any LIMS system is the balance between complete control over data tags/items to be collected from users, vs complete flexibility where software is intelligent enough to handle what a user is providing or requesting. Designing too much control makes the utility “brittle” and incapable of handling slight deviations from the original data management pipeline, posing challenges to users who want to use the LIMS system but are frustrated by strict data management policies imposed by the structure of the system. On the other hand, designing a highly flexible system that is sufficiently light-weight for general distribution is very challenging.

LIMS or data curation software employed across the NMR data ecosystem can be organized into three main groups. First, there are LIMS that seek to archive and track sample production. Examples include SPINE^22^, ProteinTracker^26^, Sesame^27^, and PiMS^28^ to name a few. Across these sample production specific LIMS, the schemas are quite different from each other as they serve different needs, processes, and communities..

Second, are data/software communities and packages that organize the software needed to record and process NMR data, and to track intermediate and final results of a data analysis pipeline. Examples of these include SPINS^24,25^, CCPN^31^, NMRFAM-SPARKY^29^ and NMRbox^30^. The applications and packages that make up this second set are not databases that store FIDs or processed NMR spectra in a relational database. They represent software suites where software conventions, versions, and data input and output formats are standardized.

The third group of data organization and curation software in the NMR field is the global community standards for making NMR data and structures publicly available. The BMRB is the main public data repository for magnetic resonance data types and molecular structures. The BMRB schema also organizes sample details, spectrometer and probe information, pulse sequence and experimental details, and is the international archive for many different NMR data types. The BMRB schema also has a textual-based archive format called NMR **<u>S</u>**tandard **<u>T</u>**ext **<u>A</u>**rchival and **<u>R</u>**etrieval (NMR-STAR) format^33^. Using NMR-STAR, NMR experiments can be recorded in a text based, machine-readable format for deposition to the BMRB, as well as storage of NMR data and experiments in a standard, well-defined ontology. Alongside NMR-STAR is the **<u>N</u>**MR **<u>E</u>**xchange **<u>F</u>**ormat^37^ (NEF), a different textual ontology to describe NMR experiments and data. NEF has particular value as a light-weight NMR restraint exchange format. NMR-STAR and NEF are standard ontologies and schema to archive and/or share NMR data and experiment descriptions, but they are not databases designed to save reproducible descriptions of NMR experiments and the collected FIDs from an experiment. Researchers will typically utilize NEF only well into a NMR study (e.g. for structural modeling) and interact with the BMRB only at a late stage of a project, after most of the study has been completed.

SpecDB is designed to allow archiving of NMR FID data immediately after data collection at the spectrometer. It does not handle any stage of post-processing of NMR FIDs, so SpecDB does not fit into the second grouping of software packages for NMR analysis described above. NMR FID data are an important data resource that will serve as input into future data mining and machine learning efforts. SpecDB fulfills the timely need for a light-weight database that can reliably organize NMR experiments as they are being collected, where the raw FID data is the central data item in the database along with experiment and sample metadata. SpecDB also supports data interchange into other FID deposition formats, like NMR-STAR.

The FID as a data item in the SpecDB schema represents a significant shift in the understanding of LIMS for NMR data. Historically, FID binary files presented a challenge for digital storage due to their size and limits on available storage. Dedicated servers or archival media (e.g.removable disks, tapes) are usually used to store FIDs. Although a separate database might be available to organize the metadata for the experiment, in most cases the connection between the FID data and the sample metadata is provided only through a physical laboratory notebook. In some LIMS systems, FID datasets are accessed through a filesystem path designating where the FID is located on the filesystem or archival media. For instance, in SPINE and SPINS, NMR data was recorded and tracked, but the FIDs sit in hierarchical directories linked to these metadata via filesystem paths. SPINE stores a wide range of experimental data and valuable information, yet the raw NMR experimental data is outside the relational nature of SPINE, leaving it vulnerable to separation from the metadata, data loss, and security issues. Presently, storage and memory resource limitations are not as much of a concern as they were a few decades ago, and relational databases can directly archive several hundred or thousands of binary files from multidimensional NMR experiments. Storing FIDs directly into the database also protects against data loss as the FIDs are internal to the database and associated with their metadata descriptions. SpecDB provides storage of FID data directly into a relational database as data items themselves.

SpecDB does not make an effort at this time to store processed frequency-domain NMR spectra. Since processed spectra files are much larger than the FID data from which they are generated, they present larger memory and storage challenges. However, it is possible to archive processing scripts in SpecDB (e.g. NMRPipe^38^ processing scripts), allowing regeneration of specific frequency-domain processed spectra. It would also be useful to have a database of such processed spectra (or scripts), prepared by NMR processing experts, for machine learning applications, but this is beyond the scope of the current version of SpecDB.

Using the Structured Query Language (SQL) to construct a relational database allows for structured queries to be completed by the NMR experimentalist, and ultimately data scientists analyzing these data post data collection. In the biomolecular NMR community FID data (as well as processed spectra) are typically stored in a file system. These data are often left to be organized by the specific researcher for a particular project. The standards/conventions employed by one researcher to organize their data collection may not be consistent across the community, or even within a research team. For example, if it were necessary to collect FID data generated in a specific date range without a database, relational or otherwise, custom software would be needed to accomplish the task. These issues are addressed by SpecDB, which provides a uniform and query-able platform for organizing NMR FID data within a research laboratory or NMR facility, and a path to sharing these data across the scientific community.

SpecDB uses JSON files to record metadata information about an NMR experiment. Data items in JSON files can be used in SQL queries. When an FID is collected at the NMR spectrometer system, other auxiliary files are also created by the data collection software, including data collection parameter files, the pulse sequence mini-program, and various spectrometer-specific acquisition files including waveform and shim files. These auxiliary files are critical to allow reproducibility of an NMR experiment. For this reason, the entire data collection directory needs to be captured and stored in the SpecDB database. To allow for queries on these data items in spectrometer files, some of them, such as the date(s) when the data were collected, and the temperature of data collection, are automatically pulled from the data collection files into JSON files, and then archived in tables of the relational database. Future query requirements can be supported by adding additional data items (e.g. NOESY mixing time) to the set of items pulled from the data collection parameter files and supported by the JSON files and SpecDB.

The command line tool of SpecDB has features similar to *git*, the command line tool to manage software projects involving many developers. In *git*, there are subcommands like *status, add, commit*, etc that are all particular steps in the tracking and maintaining a software codebase with many collaborators. In *git*, new files are added and committed to the repository at the discretion of the developer. Similar to *git*, the NMR experimentalist inserts FIDs into a SpecDB database when they determine they have recorded a complete set of metadata items for the session in the JSON file. In essence, SpecDB command line tools track the status of JSON files that contain the same data items and tables as the relational schema, and insert all the corresponding information correctly into the database.

JSON files also allow for innovative approaches for harvesting metadata for SpecDB. The initial distribution of SpecDB includes tools for using Google Sheets for metadata entry and conversion to JSON files. In our own lab, we are also exploring Microsoft Excel files, WordPress forms, and the commercial LabArchive electronic laboratory notebook tools for this purpose. We intentionally reserve judgment on recommending the “best solution” to this data harvest problem, since this will be laboratory dependent. The input to SpecDB is, ultimately, the JSON files, and various approaches can be taken to create these files.

SpecDB provides a lightweight, flexible, and robust schema and tools to archive time domain data of NMR experiments. As mentioned in the Introduction, there is a community-wide effort to expand the manner and standards for NMR researchers to deposit the raw FIDs that support their studies. Deposition of time-domain data is a major recommendation and goal of the wwPDB NMR Validation Task force for rigor and reproducibility in biomolecular NMR studies. The BMRB is collecting time-domain data, and using SpecDB able to produce NMR-STAR files for an FID is an important next step for wide-adoption of policies and practices for deposition of raw FID data.

## Conclusion

The goal of SpecDB was to build a relational schema and software to collect and track data items about biomolecular NMR samples and FID data sets, with the primary purpose of archiving and sharing these NMR time domain data. Standardized approaches for archiving FID data in relational databases provides the opportunity to develop rich datasets needed to learn new approaches for NMR data analysis. Although developed primarily using solution NMR data for proteins and nucleic acids recorded on Bruker NMR spectrometer systems, SpecDB can be easily generalized for archiving also solid-state NMR data, NMR data for oligosaccharides or small molecules, and data obtained on Varian, Agilent, JOEL, or Q-One NMR spectrometer systems. Broad use of SpecDB has the potential to create a rich data resource for a wide range of machine learning applications for biomolecular NMR. SpecDB is publicly available under the MIT open source license at the following GitHub repository: https://github.rpi.edu/RPIBioinformatics/SpecDB. The repository comes with a installation instructions and tutorials to get started with SpecDB.

## Author Contributions

All authors contributed to the design of the database schema. KF wrote the SpecDB database code. The manuscript was written with contributions from all authors.

## Acknowledgments

We thank Drs. S. Aviran, E. Baldwin, J. Hoch and J. Wedell for helpful discussions and advice.

## Supplementary Material

**Table S1:** Controlled vocabulary for the allowed tube types in SpecDB. The table above lists the allowed tube types for SpecDB. If a user attempts to indicate a tube type different from the tube name in this list, then insertion into the database will be prevented and the problem logged into the SpecDB log file. Users can edit the controlled vocabulary for the tube names in their own SpecDB instances by amending the Tubes table.

|  |  |
| --- | --- |
| Allowed NMR<br>tube type names | "0.5 mL Eppendorf" |
|  | "1.7 mL Eppendorf" |
|  | "15 mL cent. tube" |
|  | "50 mL cent. tube" |
|  | "1 L cent. tube" |
|  | "8-well PCR strip" |
|  | "NMR tube" |
|  | "plate well" |
|  | "1-mm NMR tube" |
|  | "1.7-mm NMR tube" |
|  | "3-mm NMR tube" |
|  | "3-mm Shigemi tube" |
|  | "4-mm NMR tube" |
|  | "4-mm Shigemi tube" |
|  | "5-mm NMR tube" |
|  | "5-mm Shigemi tube" |
|  | "8-mm NMR tube" |
|  | "8-mm Shigemi tube" |
|  | "10-mm NMR tube" |
|  | "10-mm Shigemi tube" |

**Table S2:** Controlled vocabulary for different isotope labeling methods. The isotopic labeling terms are data types described in the Purification Batch table. Protein samples can contain various isotope labeling schemes, and the terms above captures several common isotope labeling schemes. For isotope labeling that does not fit in these isotope labeling terms listed above, there is a separate *isotope_labeling_remark* in the Purification Batch table supporting isotope labeling methods not modeled across these columns.

| SpecDB data type | Description | Controlled Vocabulary Description |
| --- | --- | --- |
| *iso_13c_enrichment* | Describe the Carbon-13 isotopic enrichment | Must have substring ( " % 13C " ) |
| *iso_15n_enrichment* | Describe the Nitrogen-15 isotopic enrichment | Must have substring ( " % 15N " ) |
| *iso_2h_enrichment* | Describe the Deuterium isotopic enrichment | Must have substring ( " % 2H " ) |
| *iso_19f_Trp_enrichment* | Describe the Fluorine-19 isotopic enrichment on Tryptophan amino acids | Must have substring ( " % 19F-Trp " ) |
| *iso_19f_Phe_enrichment* | Describe the Fluorine-19 isotopic enrichment on Phenylalanine amino acids | Must have substring ( " % 19F-Phe " ) |
| *iso_1hd1_Leu_methyl_enrichment* | Describe the Deuterium isotopic enrichment for Leu-HD1 protons | Must have substring ( " % 1HD1-Leu " ) |
| *iso_1hd2_Leu_methyl_enrichment* | Describe the Deuterium isotopic enrichment for Leu-HD2 protons | Must have substring ( " % 1HD2-Leu " ) |
| *iso_1hd_Ile_methyl_enrichment* | Describe the Deuterium isotopic enrichment for Ile-HD protons | Must have substring ( " % 1HD-Ile " ) |
| *iso_1hg1_Val_methyl_enrichment* | Describe the Deuterium isotopic enrichment for Val-HG1 protons | Must have substring ( " % 1HG1-Val " ) |
| *iso_1hg2_Val_methyl_enrichment* | Describe the Deuterium isotopic enrichment for Val-HG2 protons | Must have substring ( " % 1HG2-Val " ) |
| *iso_1hb_Ala_methyl_enrichment* | Describe the Deuterium isotopic enrichment for Ile-HD protons | Must have substring ( " % 1HD-Ile " ) |

